# Electrostatic Plasma Membrane Targeting Contributes to Dlg Function in Cell Polarity and Tumorigenesis

**DOI:** 10.1101/2020.10.08.331603

**Authors:** Juan Lu, Wei Dong, Yan Tao, Yang Hong

## Abstract

Discs large (Dlg) is an essential polarity protein and a tumor suppressor originally characterized in *Drosophila* but is also well conserved in vertebrates. Like the majority of polarity proteins, plasma membrane (PM)/cortical localization of Dlg is required for its function in polarity and tumorigenesis, but the exact mechanisms targeting Dlg to PM remain to be fully elucidated. Here we show that, similar to the recently discovered polybasic polarity proteins such as Lgl and aPKC, Dlg also contains a positively charged polybasic domain that electrostatically binds the PM phosphoinositides PI4P and PI(4,5)P_2_. Electrostatic targeting by the polybasic domain contributes significantly to the PM localization of Dlg in follicular and early embryonic epithelial cells, and is crucial for Dlg to regulate both polarity and tumorigenesis. The electrostatic PM targeting of Dlg is controlled by a potential phosphorylation-dependent allosteric regulation of its polybasic domain, and is specifically enhanced by the interactions between Dlg and another basolateral polarity protein and tumor suppressor Scrib. Our studies highlight an increasingly significant role of electrostatic PM targeting of polarity proteins in regulating cell polarity.

## INTRODUCTION

Polarity proteins play conserved and essential roles in regulating the apical-basal polarity in epithelial cells of both invertebrates and vertebrates. Among them, Discs large (Dlg), Scrib and Lgl also act as tumor suppressors and share the same basolateral subcellular localization in epithelial cells (Bilder et al., 2003; Tanentzapf and Tepass, 2003). Like many polarity proteins, the plasma membrane (PM)/cortical localization is essential for Dlg, Lgl and Scrib for regulating apical-basal polarity and tumorigenesis (Hough et al., 1997; Ventura et al., 2020). Recent studies discovered that multiple polarity proteins contain so-called polybasic (PB) domains that are typically of 20-40aa length and are highly positively charged due to enriched Arg and Lys residues (Hammond and Hong, 2017). Polybasic (domain) polarity proteins like Lgl and aPKC can specifically target to PM by electrostatically binding the negatively charged phospholipids, in particular PI4P and PI(4,5)P_2_ (PIP2) whose unique enrichment in PM makes the inner surface of PM the most negatively charged in the cell (Hammond et al., 2012). While the electrostatic PM targeting of Lgl is now well established (Bailey and Prehoda, 2015; Dong et al., 2015), the exact mechanisms targeting Dlg to PM remain to be fully elucidated. Here we report that Dlg, like recently characterized polybasic polarity proteins Lgl and aPKC (Dong et al., 2020), contains a polybasic domain that electrostatically targets Dlg to the PM and is necessary for Dlg to regulate polarity and tumorigenesis. Our results also suggest that Scrib specifically enhances the electrostatic PM targeting of Dlg through regulating a potential phosphorylation-dependent allosteric control mechanism.

## RESULTS AND DISCUSSION

### A polybasic domain in Dlg mediates its PM targeting

Dlg belongs to the MAGUK protein family and contains domains of PDZ, SH3, HOOK and guanylate kinase (GUK) (Fig. 1A). The GUK domain is considered kinase dead and instead acts as protein interacting domain that can bind SH3 domain either intramolecullarly or intermolecullarly (McGee et al., 2001) (see below). By sequence analysis we identified a well conserved candidate polybasic domain that spans the C-terminal half of SH3 domain and the N-terminal half of HOOK domain (Fig. 1A), with a basic-hydrophobic index (Brzeska et al., 2010) of 0.90, comparable to polybasic domains in Lgl (1.01), aPKC (0.96), and Numb (1.07). In liposome pull-down assays, GST fusion of Dlg polybasic domain (GST-PB) bound both PI4P- and PIP2-liposomes but not liposomes containing only phosphatidylserine (PS) (Fig. 1A). GST-PB-KR6Q or GST-PB-KR15Q in which positive charges were either partially or completely eliminated by K/R->Q mutations did not bind either liposomes (Fig. 1A), supporting an electrostatic interaction between the positively charged polybasic domain and the negatively charged PI4P- or PIP2-membrane.

**Figure 1.**
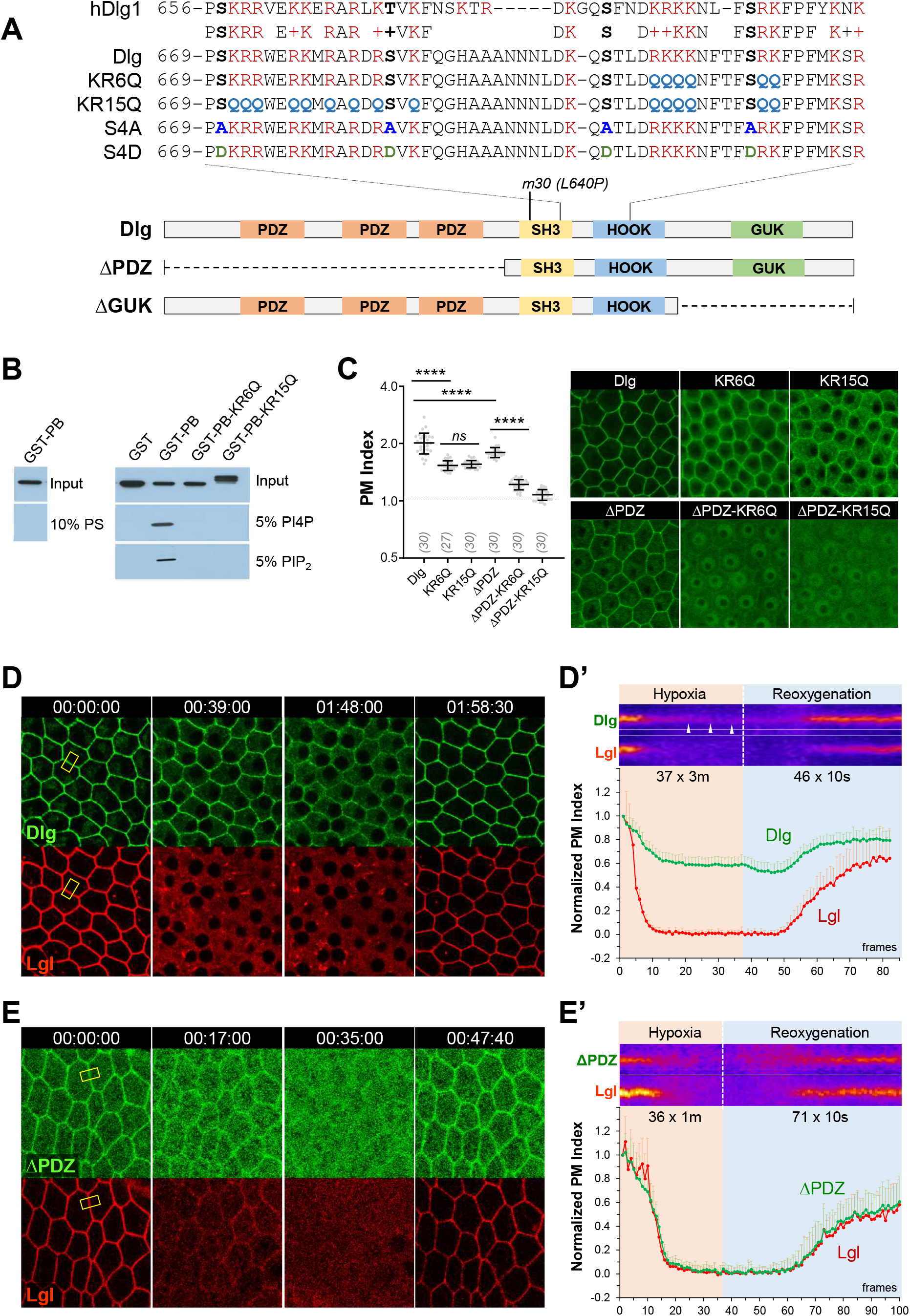
A polybasic domain in Dlg mediates its electrostatic PM targeting. (**A**) Polybasic domains in human Dlg1 (hDlg1, NP_004078.2) and *Drosophila* Dlg (NP_996405). Mutations in Dlg^KR6Q^, Dlg^KR15Q^, Dlg^S4A^ and Dlg^S4D^, as well as the point mutation of *dlg*^*m30*^ (“*m30*”) allele, are also shown. Bottom: deletions in Dlg^ΔPDZ^ and DlgΔGUK. (**B**) Western blot by GST antibody showing that GST-PB, but not GST or GST-PB-KR6Q or GST-PB-KR15Q, co-sedimented with PI4P- and PIP2-liposomes. (**C**) PM localization of wild type and mutant Dlg::GFP in follicular cells. PM Index: values above 1 (dashed line) indicate predominant PM localization while below 1 indicate cytosolic localization. In parentheses: sample numbers (*n*). (**D, E**) Follicular cells expressing Dlg::GFP^KI^ and Lgl::mCherry (D, Movie S1), or Dlg^ΔPDZ^::GFP and Lgl::mCherry (E, Movie S3), undergoing hypoxia followed by reoxygenation. Kymographs were sampled at the boxes indicated in D and E. Arrowheads in kymographs highlight the persistent residual PM localization of Dlg under hypoxia. (**D’, E’**) Quantification of Dlg and Lgl PM localization (*n*=20, 20). Time stamps in *hh:mm:ss* format. ****: p<0.00001. *ns*: p>0.05.

To investigate whether the polybasic domain is required for targeting Dlg to PM in vivo, we generated transgenic flies expressing Dlg::GFP, Dlg^KR6Q^::GFP and Dlg^KR15Q^::GFP under the ubiquitin promoter (Dong et al., 2020). Although *dlg* locus apparently expresses numerous isoforms, our *ubi-dlg::GFP* (“*dlg::GFP*”) based on isoform-G fully rescued the null mutants of *dlg*^*[A]*^ and expressed at levels similar to an endogenous GFP knock-in allele (“*dlg::GFP*^*KI*^”) (Fig. S1A). Dlg::GFP showed typical basolateral PM/cortical localization in follicular and embryonic epithelial cells (Fig. 1C, S2A), while both Dlg^KR6Q^ and Dlg^KR15Q^ showed strong reduction of PM localization (Fig. 1C, S2C) and failed to rescue *dlg*^*[A]*^ (Fig. 4D). Dlg interacts with multiple proteins on the PM through its three PDZ domains (Hough et al., 1997; Qian and Prehoda, 2006), and Dlg^ΔPDZ^::GFP also showed reduced PM localization in follicular cells, albeit less severe than Dlg^KR6Q^ and Dlg^KR15Q^ (Fig. 1C). Consistent with that both PDZ and polybasic domains contribute to PM targeting in follicular cells, Dlg^ΔPDZ-KR6Q^::GFP and Dlg^ΔPDZ-KR15Q^::GFP became virtually lost from PM (Fig. 1C).

To further investigate the electrostatic PM targeting of Dlg in vivo, we used a previously established assay that applies controlled hypoxia to acutely and reversibly deplete PIP2 and PI4P in the PM (Dong et al., 2015) (data not shown). Similar to polybasic polarity proteins Lgl (Dong et al., 2015) and aPKC (Dong et al., 2020), hypoxia also induced acute and reversible loss of Dlg::GFP^KI^, as well as ubi-Dlg::GFP, from PM in follicular cells (Fig. 1D, Movies S1, S2). However, unlike Lgl which became completely cytosolic within 30-60min of hypoxia, residual Dlg::GFP^KI^ persisted on PM over 100min of hypoxia (Fig. 1D, S1B). In contrast, hypoxia induced complete PM loss of Dlg^ΔPDZ^::GFP that is identical to Lgl (Fig. 1E, S1B, Movie S3). Such data suggest that in follicular cells a significant portion of Dlg electrostatically binds to PM, while the rest were PM-retained by PDZ domain-dependent interactions.

Dlg is also a component of septate junction (SJ) which is composed of over twenty different proteins (Izumi and Furuse, 2014), making SJ the primary target of PDZ-dependent PM localization. In early embryonic epithelia cells with undeveloped SJ, Dlg::GFP was readily lost from PM under hypoxia, suggesting that its PM targeting is mostly electrostatic (Fig. S2A). In stage 14 or later embryos with mature SJ, PM localization of Dlg::GFP^KI^ was significantly enhanced (Fig. S2C, D) and became completely resistant to hypoxia (Fig. S2A). In contrast, in stage 14 embryos mutant of core SJ proteins such as Coracle (*cora*) (Lamb et al., 1998) or Na^+^,K^+^-ATPase α-subunit (*Atp*α) (Paul et al., 2003), Dlg::GFP^KI^ PM localization became sensitive to hypoxia (Fig. S2B). In addition, enhanced PM localization in late embryos was seen in electrostatic Dlg mutants, but not in mutants lacking PDZ domains (Fig. S2C, D), consistent with that PDZ domains are required for Dlg localization to SJ (Hough et al., 1997). Indeed, PM localization of Dlg^ΔPDZ^::GFP in late embryos is electrostatic, based on its sensitivity to hypoxia (Fig. S2B).

In summary, with the help of hypoxia-based imaging assays we were able to specifically probe the electrostatic PM targeting of Dlg in vivo. Our data revealed that in follicular and early embryonic epithelial cells full PM localization of Dlg requires both electrostatic PM targeting by the polybasic domain and protein-interactions mediated by PDZ domains, while in epithelial cells with mature septate junctions PDZ domains alone can act redundantly to retain Dlg on the PM.

### PM targeting of Dlg depends on PI4P and PIP2 in vivo

PM PI4P and PIP2 are majorly responsible for electrostatically binding polybasic proteins (Hammond, 2012). To investigate the role of PIP2 in targeting Dlg in vivo, we acutely depleted PIP2 in follicular cells, using an inducible system (Dong et al., 2020; Dong et al., 2015; Reversi et al., 2014) in which rapamycin addition induces dimerization between a FRB-tagged PM anchor protein (i.e. Lck-FRB-CFP) and a FKBP-tagged phosphatase (i.e. RFP-FKBP-INPP5E), resulting in acute PM recruitment of INPP5E that rapidly depletes PM PIP2. For reasons unknown, PM levels of Dlg::GFP^KI^ or Dlg^ΔPDZ^::GFP increased in cells expressing Lck-FRB and FKBP-INPP5E, regardless of DMSO or rapamycin treatment (Fig. 2A). Nonetheless, under rapamycin but not DMSO treatment, PM localization of both Dlg::GFP^KI^ and Dlg^ΔPDZ^::GFP showed strong reduction in cells expressing Lck-FRB and FKBP-INPP5E (Fig. 2A), supporting that Dlg PM targeting is at least partially dependent on PM PIP2.

**Figure 2.**
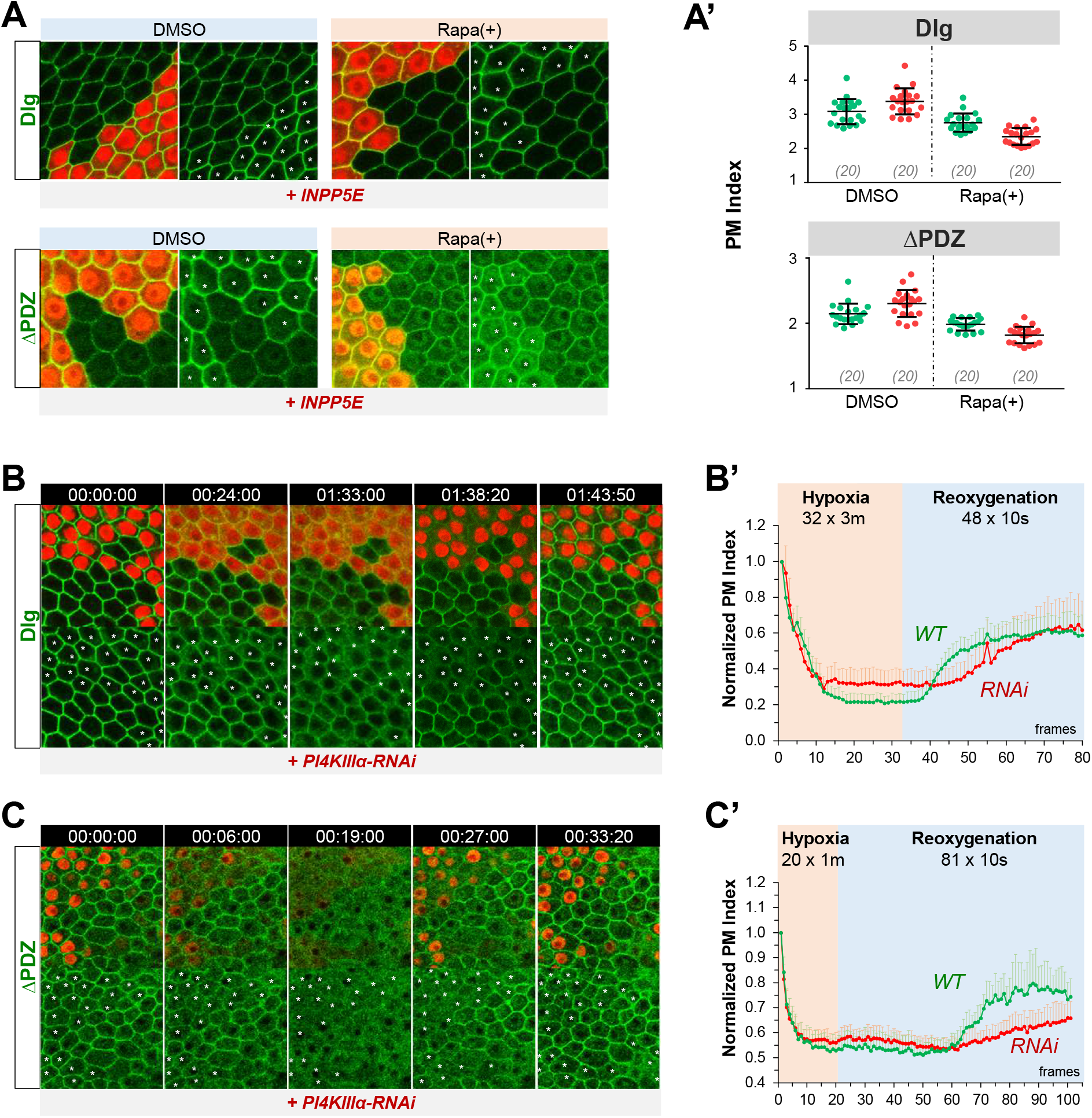
Electrostatic PM targeting of Dlg depends on PI4P and PIP2. (**A**) *dlg*^*KI*^*::GFP* or *dlg*Δ*PDZ::GFP* follicular epithelia were treated with either DMSO (control) or rapamycin (“rapa(+)”) and imaged live. Cells expressing Lck-FRB-CFP (not imaged) and RFP-FKBP-INPP5E (“INPP5E”) were labeled by nuclear RFP. (**A’**) Quantification of PM localization of Dlg::GFP or Dlg^ΔPDZ^::GFP in wild type (green dots) and INPP5E (red dots) cells. In parentheses: sample numbers. (**B, C**) Follicular cells expressing Dlg^KI^::GFP (B, Movie S4) or Dlg^ΔPDZ^::GFP (C, Movie S5) undergoing hypoxia and reoxygenation. *PI4KIII*α*-RNAi* cells were labeled by RFP. (**B’, C’**) Quantification of Dlg::GFP (B’) or Dlg^ΔPDZ^::GFP (C’) PM localization in both wild type (green lines) and *PI4KIII*α*-RNAi* (red lines) cells (*n*=20, 20).

At present, similar tools for acutely depleting PI4P or both PI4P and PIP2 are not available in *Drosophila*. We thus manipulated PI4P levels genetically by RNAi-knock down of PI4KIIIα, the PtdIns-4-kinase specifically responsible for maintaining the PM PI4P levels (Bojjireddy et al., 2014; Tan et al., 2014). Wild type cells and *PI4KIII*α*-RNAi* cells showed similar PM levels of Dlg::GFP^KI^ and Dlg^ΔPDZ^::GFP and rates of PM delocalization under hypoxia (Fig. 2B, Movies S4, S5). However, consistent with that PI4KIIIα is likely necessary for replenishing PM PI4P and PIP2 (Bojjireddy et al., 2014), PM recoveries of Dlg::GFP^KI^ and Dlg^ΔPDZ^::GFP were significantly delayed in *PI4KIII*α*-RNAi* cells (Fig. 2B). Overall, our data suggest that PI4P and PIP2 act redundantly to electrostatically bind Dlg to PM.

### Potential phosphorylation events on the polybasic domain regulate the PM targeting of Dlg

Polybasic domains are often regulated by direct phosphorylations which inhibit its PM binding by neutralizing the positive charges (Hong, 2018), such as in the cases of Lgl, Numb and Miranda (Bailey and Prehoda, 2015; Dong et al., 2015). To investigate whether Dlg polybasic domain is also regulated by phosphorylation events, we mutated four well conserved S/T sites within its polybasic domain (Fig. 1A), of which the first two are the previously characterized phosphorylation sites of aPKC and PKCα (Golub et al., 2017; O’Neill et al., 2011). Both phosphomemitic Dlg^S4D^::GFP and non-phosphorylatable Dlg^S4A^::GFP showed significant loss from PM (Fig. 3A), suggesting that these potential phosphorylation events unlikely regulate the polybasic domain through charge neutralization. Interestingly, PM localization of Dlg^S4A^::GFP, albeit reduced, became resistant to hypoxia (Fig. S3A, Movie S6), while removing PDZ domains in Dlg^S4A^::GFP nearly abolished the PM localization of Dlg^ΔPDZ-S4A^::GFP (Fig. 3A), suggesting that the residual PM localization of Dlg^S4A^::GFP is non-electrostatic but PDZ-dependent.

**Figure 3.**
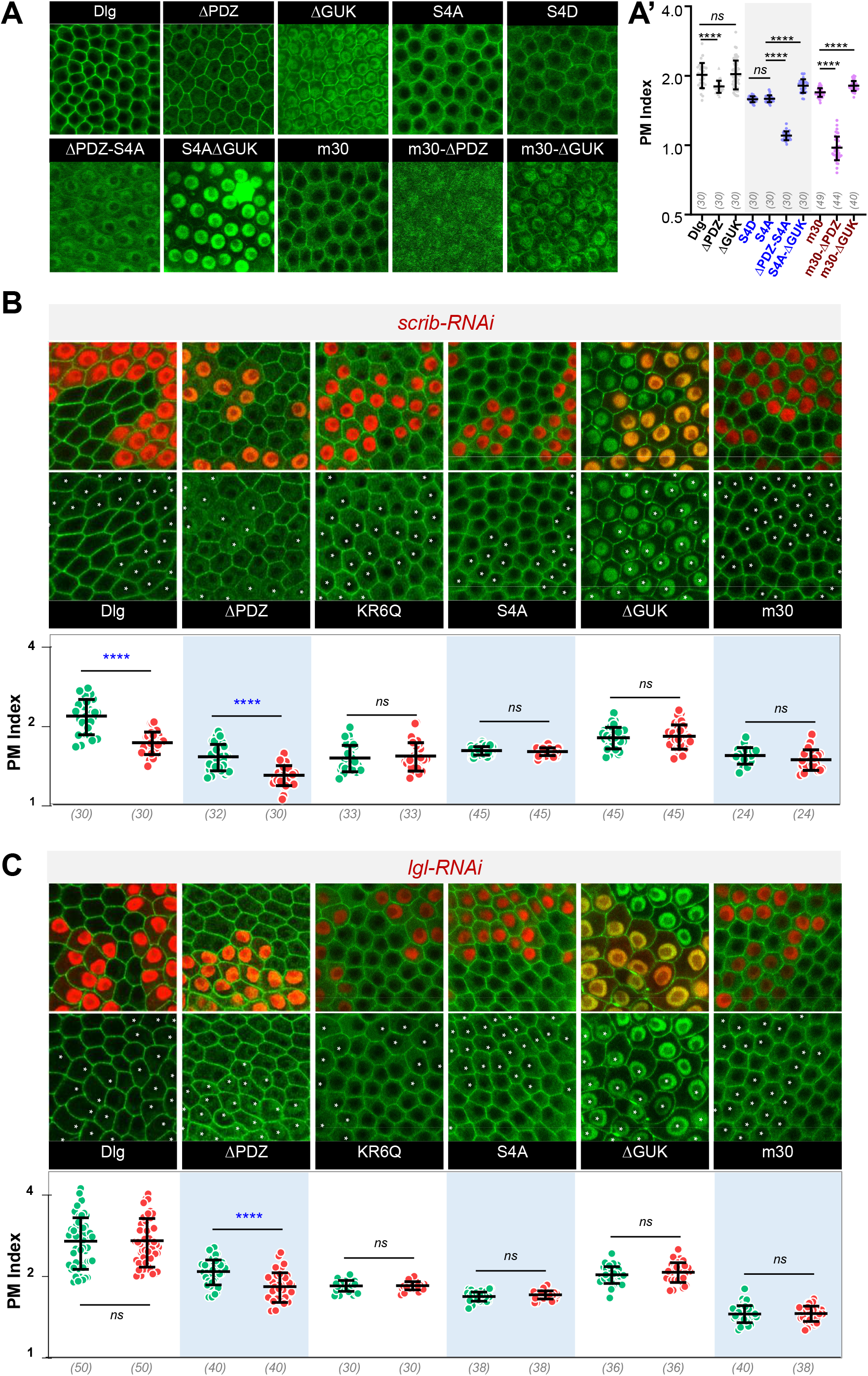
Scrib enhances the electrostatic PM targeting of Dlg. (**A**) Follicular cells expressing the Dlg::GFP or mutants as indicated. PM localization of each mutant was quantified in A’. (**B, C**) Follicular cells expressing Dlg::GFP or mutants as indicated. *scrib-RNAi* (**B**) or *lgl-RNAi* (**C**) cells were labeled by RFP. (**B’, C’**) Quantifications of wild type and mutant Dlg::GFP PM localizations in wild type (green dots) and *RNAi* (red dots) cells.

Our recent studies also showed that the polybasic domain in aPKC, which is the pseudosubstrate region (PSr) of no characterized phosphorylation sites, is allosterically regulated (Dong et al., 2020). The PSr in aPKC binds and autoinhibits the kinase domain which in turn occludes PSr from electrostatically binding to PM, while binding of Par-6 to aPKC induces conformation changes exposing the PSr to PM-binding. Given that SH3 and GUK domains in Dlg could also bind each other, we postulate that the loss of potential phosphorylation events in Dlg^S4A^::GFP keeps the SH3-GUK in a closed conformation that occludes the polybasic domain from binding to PM. Removing the GUK domain however made DlgΔGUK::GFP localize both on PM and in nuclei (Fig. 3A), likely due to that the Arg/Lyn-rich feature of polybasic domain is also similar to nuclear localization signal (NLS) (Dong et al., 2015). Consistent with that removing the GUK domain exposes the polybasic domain, Dlg^ΔGUK^::GFP PM localization is partially electrostatic just like Dlg::GFP (Fig. S2B, Movie S7). In addition, Dlg^ΔGUK^::GFP’s moderate nuclear localization can be overcome by its interaction with SJ, as Dlg^ΔGUK^::GFP was SJ-localized in larval wing disc epithelia which develop strong SJ (Fig. S2E) (Hough et al., 1997). In contrast, Dlg^S4A-ΔGUK^::GFP showed overwhelmingly high nuclear localization both in follicular cells (Fig. 3A) and in wing imaginal epithelial cells (Fig. S2E), suggesting that removing the GUK likely also exposes the polybasic domain in Dlg^S4A^, but the lack of potential phosphorylation events strongly enhanced the nuclear localization property of the polybasic domain.

Although additional studies are needed to confirm the phosphorylation events of Dlg in vivo, our results are consistent with a model that the polybasic domain in Dlg is regulated by phosphorylation events that allosterically relieve its blockage by GUK and promote its PM targeting over nuclear localization (Fig. 4E).

**Figure 4.**
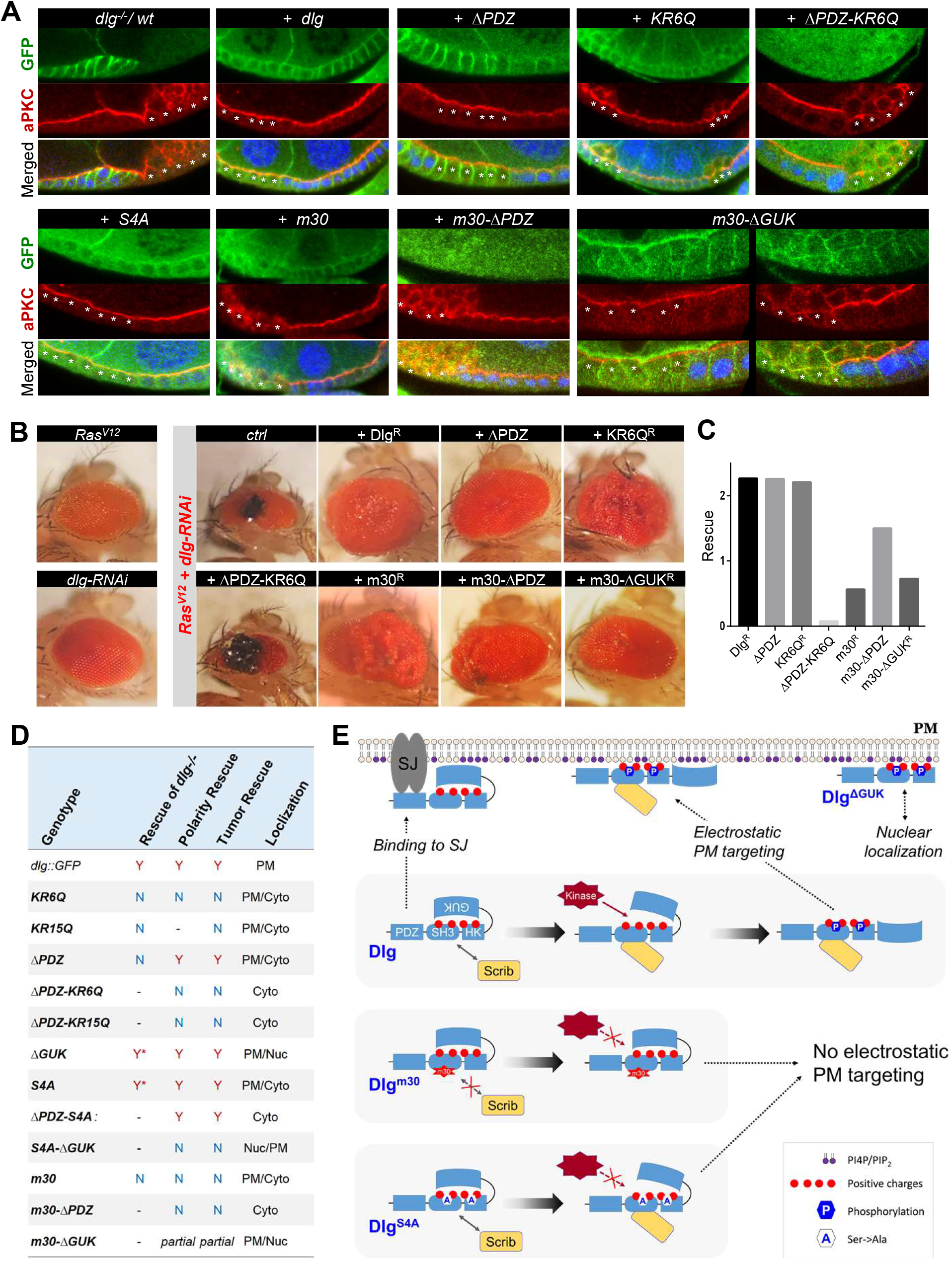
Electrostatic PM targeting of Dlg regulates cell polarity and tumorigenesis. (**A**) *dlg*^*-/-*^ follicular cells were marked by the loss of RFP (blue in all merged images), except for “*dlg*^*-/-*^ *wt*” in which the *dlg*^*-/-*^ clones were marked by the absence of Dlg (green). Samples were expressing wild type or mutant Dlg::GFP as indicated, and stained for GFP (green) and aPKC (red). Note that *dlg*^*-/-*^, *dlg*^*m30-*^Δ*GUK::GFP* clones showed roughly equal frequency of rescued and non-rescued polarity defects. Asterisks highlight *dlg*^*-/-*^ mutant cells. (**B**) Eyes from adult flies of *Ras*^*V12*^, or *dlg-RNAi* or *Ras*^*V12*^*/dlg-RNAi* combined with additional expression of wild type and mutant Dlg as indicated. “*ctrl*”: *Ras*^*V12*^*/dlg-RNAi* only. (**C**) Quantifications of the rescue of *Ras*^*V12*^*/dlg-RNAi* lethality by wild type and mutant Dlg as indicated. (**D**) Summary of genetic rescue analyses of *ubi-dlg::GFP* transgenic alleles. *: sterile; Nuc: nuclear; Cyto: cytosolic; PM: plasma membrane; -: not done. (**E**) A hypothetical model on the potential phosphorylation-dependent allosteric control of Dlg polybasic domain and its regulation by Scrib. Potential regulatory defects in Dlg^m30^, Dlg^S4A^ and DlgΔUGK are also shown. Kinases phosphorylating Dlg’s polybasic domain remain to be characterized but could include aPKC and PKCα. The sequential order between Scrib-binding to Dlg and its phosphorylation is postulated. SJ: septate junctions.

### Scrib specifically enhances the electrostatic PM targeting of Dlg

Scrib and Lgl colocalize with Dlg at basolateral PM and play similar functions in cell polarity and tumorigenesis, but it is unknown how Scrib and Lgl may specifically regulate the electrostatic PM targeting of Dlg. Consistent with recent studies that Scrib but not Lgl appears to physically interact with Dlg (Khoury and Bilder, 2020; Ventura et al., 2020), knocking down Scrib by RNAi induced a partial loss of Dlg from PM (Fig. 3B). More importantly, Scrib appears to specifically enhance the electrostatic PM targeting of Dlg, as Dlg^ΔPDZ^, which can only bind PM electrostatically, was dramatically reduced from PM in *scrib-RNAi* cells (Fig. 3B). In contrast, the residual PM localization of electrostatic mutant Dlg^KR6Q^ was not affected by *scrib-RNAi* (Fig. 3B). Scrib interaction depends on the SH3 domain in Dlg and a single point mutation in SH3 domain in *dlg*^*m30*^ mutant (Fig.1A) abolishes such interaction (Khoury and Bilder, 2020). Indeed, similar to electrostatic mutants such as Dlg^KR6Q^, Dlg^S4A^ and Dlg^ΔGUK^, PM localization of Dlg^m30^::GFP was also reduced and became resistant to *scrib-RNAi* (Fig. 3B), supporting that interactions between Scrib and Dlg are necessary for Scrib to enhance Dlg electrostatic PM targeting. More importantly, PM localization of Dlg^S4A^ or Dlg^ΔGUK^, two mutants defective in the potential phosphorylation-dependent allosteric control of polybasic domain, was also resistant to *scrib-RNAi* (Fig. 3B), suggesting that Scrib may act by facilitating such phosphorylation-allosteric mechanism (Fig. 4E).

In agreement with the recent findings that Lgl does not appear to physically associate with Dlg and Scrib (Khoury and Bilder, 2020; Ventura et al., 2020), *lgl-RNAi* did not affect the PM localization of Dlg, Dlg^KR6Q^, Dlg^S4A^, Dlg^m30^ and Dlg^ΔGUK^ (Fig. 3C, S4C””). However, PM localization of Dlg^ΔPDZ^ was mildly reduced in *lgl-RNAi* cells (Fig. 3C). Thus Lgl could also play a moderate role in enhancing Dlg electrostatic PM targeting, through mechanisms presently unclear. Finally, knocking down GukH, which interacts with the GUK domain and is required for aPKC-regulation of Dlg in neuroblasts (Golub et al., 2017), did not affect the electrostatic PM targeting of Dlg^ΔPDZ^ (Fig. S4C’), consistent with that GukH appears to be dispensable for epithelial polarity (Caria et al., 2018).

### Electrostatic PM targeting is essential for Dlg to regulate cell polarity and tumorigenesis

We also investigated how electrostatic PM targeting may specifically contribute to Dlg function in polarity and tumorigenesis. Follicular cells of *dlg*^*-/-*^ showed dramatic disruption of cell polarity evidenced by the mislocalization of apical polarity proteins such as aPKC (Fig. 4A).

Such phenotype was fully rescued by ectopic expression of Dlg and Dlg^ΔPDZ^, but not electrostatic mutant Dlg^KR6Q^ or Dlg^ΔPDZ-KR6Q^ (Fig. 4A), suggesting that electrostatic PM targeting is essential for Dlg to regulate apical-basal polarity in follicular cells. Surprisingly, *dlg*^*[A]*^*/Y; dlg*^*S4A*^*::GFP* and *dlg*^*[A]*^*/Y*; *dlg*^*ΔGUK*^*::GFP* males are viable although sterile (Fig. 4D), thus the potential mechanisms regulating the polybasic domain may be largely dispensable for Dlg function in normal development.

We then used a well-established tumor model (Brumby et al., 2011) to investigate how electrostatic PM targeting is required for Dlg’s tumor suppressor function. Overexpression of constitutively active Ras^V12^ in larval eye disc produced a moderate rough eye phenotype (Fig. 4B). While *dlg-RNAi* alone in the eye did not produce obvious phenotypes (Caria et al., 2018), combining Ras^V12^ over-expression and *dlg-RNAi* (i.e. “*Ras*^*V12*^*/dlg-RNAi*”) produced massive eye tumors which resulted in strong larval/pupal lethality, with very few adults escaped with reduced eyes size and dark tumor tissues (Fig. 4B). We made transgenic stocks that express wild type or mutant Dlg proteins from cDNA sequences modified to be resistant to *dlg-RNAi* (Fig. S4C, Table S1). Any of the ^ΔPDZ^ mutants is also resistant to *dlg-RNAi* as the RNAi targets the sequence near the N-terminal coding region. As expected, expression of RNAi-resistant *dlg::GFP* (“Dlg^R^::GFP”) fully rescued *Ras*^*V12*^*/dlg-RNAi* lethality and eye morphology (Fig. 4B). Both Dlg^ΔPDZ^ and polybasic mutant Dlg^KR6Q-R^ rescued lethality of *Ras*^*V12*^*/dlg-RNAi*, but eyes in *dlg*^*KR6Q-R*^ flies still showed strong tissue overproliferation (Fig. 4B). In contrast, Dlg^ΔPDZ-KR6Q^ failed to rescue either lethality or eye morphology and tumorigenesis (Fig. 4B). Thus, both electrostatic and PDZ-dependent PM targeting are required for Dlg tumor suppressor function and they act redundantly in rescuing the *Ras*^*V12*^*/dlg-RNAi* tumor lethality, although electrostatic PM targeting appears to be more specifically required for inhibiting the overproliferation of *Ras*^*V12*^*/dlg-RNAi* tumor cells.

Notably, Dlg^m30-ΔGUK^, but not Dlg^m30^ or Dlg^m30-ΔPDZ^, partially rescued the polarity defects in *dlg*^*-/-*^ cells (Fig. 4A). RNAi-resistant Dlg^m30-R^ only moderately rescued the lethality caused by *Ras*^*V12*^/*dlg-RNAi*, and survivors still showed strong overproliferation in eyes (Fig. 4B). Consistent with the polarity rescue results in Fig. 4A, removing GUK but not PDZ domains in Dlg^m30^ significantly enhanced the rescue of the overproliferation phenotype in *Ras*^*V12*^/*dlg-RNAi* eyes (Fig. 4B). Such data are consistent with that eliminating the potential allosteric inhibition of polybasic domain in Dlg partially compensated loss of Scrib-dependent enhancement electrostatic PM targeting. Interestingly, although both Dlg^S4A^ and Dlg^m30^ were defective in electrostatic PM targeting, Dlg^S4A^ showed strong rescue activity whereas Dlg^m30^ did not (Fig. 4D), suggesting that Scrib/Dlg interaction is critical for Dlg function besides regulating Dlg’s electrostatic PM targeting. In addition, unlike Dlg^S4A^ or Dlg^ΔGUK^, Dlg^S4A^^ΔGUK^ showed no rescue activity (Fig. 4D), likely due to that its strong nuclear concentration made protein levels at PM too low to rescue (Fig. 4D).

The electrostatic nature of Dlg PM targeting makes Dlg a new member of the polybasic polarity protein family which includes at least Lgl, aPKC, Numb and Miranda. Our results also make it increasingly clear that electrostatic PM targeting is a key molecular mechanism widely used by polarity proteins for achieving controlled subcellular localization and for regulating cell polarity. It will be of great interest for future studies to further integrate the electrostatic PM targeting mechanism into the regulatory network of polarity protein and their interacting partners, by uncovering the essential molecular mechanisms regulating this simple but elegant physical interaction between polarity proteins and PM.

## Acknowledgements

We are grateful to Drs. David Bilder, Helena Richardson, Greg Beitel, Richard Fehon, Ulli Tepass, and Stefano De Renzis for reagents and fly stocks, Dr. Simon Watkins and University of Pittsburgh Medical School Center for Biologic Imaging for generous imaging and microscopy support, Bloomington Stock Center for fly stocks, and Developmental Studies Hybridoma Bank (DSHB) for antibodies.

## Author Contributions

Conceptualization: Y.H., W.D. J.L.; Investigation: J.L, W.D., Y.T., Y.H.; Writing - Review &Editing: Y.H; Funding acquisition: Y.H.; Supervision: Y.H.

## MATERIALS AND METHODS

### Fly Stocks

Flies of carrying transgenic *ubi-dlg::GFP* or *ubi-dlg::GFP* mutant (“*ubi-dlg**::GFP*”) alleles were generated by *phiC31*-mediated integration protocol (Huang et al., 2009). *attP*^*VK00022*^ (BL#24868) stock was used to integrate *ubi-dlg:GFP* and *ubi-dlg**::GFP* constructs to the 2^nd^ chromosome. Transgenic alleles of *ubi-dlg::GFP* and *ubi-dlg**::GFP* were further recombined with *y w dlg*^*[A]*^ *FRT19A/FM7c* (BL#57086). Summary of *ubi-dlg**::GFP* alleles are in Table S1.

*cora*^*5*^ */ CyO dfdGMR-YFP and Atp*α ^*DTS2A3*^ */ TM6 dfdGMR-YFP* are gifts from Dr. Greg Beitel, (Northwestern University, USA).

*w UASp>mRFP::FKBP-5Ptase* (“FKBP-INPP5E”) and *w;* ; *UASp>Lck-FRB::CFP* are gifts from Dr. Stefano De Renzis (EMBL Heidelberg, Germany) (Reversi et al., 2014).

*ey-Gal4, UAS-Ras*^*V12*^ stocks were gift from Dr. Helena Richardson.

*w P[w[+mC]=PTT-GC]dlg1*^*[YC0005]*^ *(“dlg::GFP*^*KI*^*”*, BL#50859*), UAS-PI4KIII*α*-RNAi* (BL#35256), *UAS-scrib-RNAi* (BL#29552), *UAS-lgl-RNAi* (BL#38989), *UAS-dlg-RNAi* (II) (BL#39035), *UAS-dlg-RNAi* (III) (BL#34854) and *gukh-RNAi* (BL#42486) (Caria et al., 2018) are from Bloomington Stock Center.

*w; lgl::mCherry* knock-in and *w; lgl::GFP* knock-in stocks were previously published (Dong et al., 2015).

*Drosophila* cultures and genetic crosses are carried out at 25°C.

### Molecular cloning

To make *ubi-dlg::GFP*, ubiquitin promoter (1872bp) was PCR amplified from plasmid pWUM6 (a gift from Dr. Jeff Sekelsky, University of North Carolina at Chapel Hill) using primers 5-AGTGTC GAATTC CGCGCAGATC GCCGATGGGC and 5-CTGGAC GCGGCCGC GGTGGATTATTCTGCGGG and inserted into pGE-attB vector (Huang et al., 2009) to generate vector pGU. DNA fragments encoding Dlg::GFP was then inserted into pGU vector. More details about DNA constructs used in this report are listed in Table S1. Sequence of the Dlg isoform used in this study can be found by NCBI RefSeq ID NP_996405.1.

### Liposome pull-down assays

Liposomal binding assays were carried out as described (Kim et al., 2008). Lipid mixture of 37.5% PC (Cat#840051C), 10% PS (Cat#840032C), 37.5% PE (Cat#840021C), 10% Cholesterol (Cat#700000P) and 5% PI(4,5)P_2_ (Cat#840046X) or PI4P (Cat#840045X, all lipids were purchased from Avanti Polar Lipids Inc) was dried and resuspended to a final concentration of 1 mg/ml of total phospholipids in HEPES buffer and subjected to 30 min sonication. Formed liposomes were harvested at 16,000g for 10 min and resuspended in binding buffer (HEPES, 20 mM, 7.4, KCl 120 mM, NaCl 20mM, EGTA 1mM, MgCl 1mM BSA 1mg/ml). Approximately 0.1μg of purified protein or protein complex was mixed with 50μl of liposome suspension in each liposome-binding assay. Liposomes were pelleted at 16,000g for 10min after 15min incubation at room temperature, and were analyzed by western blot to detect co-sediment of target protein(s).

### Generation of mitotic mutant clones in *Drosophila* follicular epithelia

Mutant follicular cell clones of *dlg*^*[A]*^ were generated by the routine FLP/FRT technique. Young females were heat-shocked at 37°C for 1 hour and their ovaries were dissected 3 days later.

### Live imaging and hypoxia treatment in *Drosophila* epithelial cells

Embryos and dissected ovaries were imaged according to previously published protocol (Dong et al., 2015; Huang et al., 2011). The embryos were staged by timing and kept in 25 °C for 2 hours before imaging.

Ovaries from adult females of 2-days old were dissected in halocarbon oil (#95). Follicular cells containing over-expressing or RNAi clones were generated by heat-shocking the young females of the correct genotype at 37°C for 15-30min and ovaries were dissected 3 days later. To ensures sufficient air exchange to samples during the imaging session, dechorionated embryos or dissected ovaries were mounted in halocarbon oil on an air-permeable membrane (YSI Membrane Model #5793, YSI Inc, Yellow Springs, OH) sealed by vacuum grease on a custom-made plastic slide over a 10×10mm^2^ cut-through window. The slide was then mounted in a custom made air-tight micro chamber for live imaging under confocal microscope. Oxygen levels inside the chamber were controlled by flow of either air or custom O_2_/N_2_ gas mixture at the rate of approximately 1-5 cc/sec. Images were captured at room temperature (25°C) on an Olympus FV1000 confocal microscope (60x Uplan FL N oil objective, NA=1.3) by Olympus FV10-ASW software, or on a Nikon A1 confocal microscope (Plan Fluo 60x oil objective, NA=1.3) by NIS-Elements AR software. Images were further processed in ImageJ and Adobe Photoshop.

### Induction of mRFP::FKBP-5Ptase and Lck-FRB::CFP dimerization in live *Drosophila* follicular cells

Young females of *w UASp>mRFP::FKBP-5Ptase /dlg::GFP;;hs-FLP Act5C(FRT*.*CD2)-Gal4 UAS-RFP/UASp>Lck-FRB::CFP* or *w UASp>mRFP::FKBP-5Ptase /+; ubi-dlg*△ *PDZ::GFP/+; hs-FLP Act5C(FRT*.*CD2)-Gal4 UAS-RFP/UASp>Lck-FRB::CFP* were heat-shocked at 37°C for 15min. Ovaries were dissected 3 days later in Schneider’s medium, mounted in a drop of 20μl Schneider’s medium containing 10μM rapamycin or DMSO on a gas-permeable slide, and imaged live as previously described (Dong et al., 2015; Huang et al., 2011).

### Immunostaining and confocal imaging

Immunostaining of follicular cells and embryos were carried out as previously described (Huang et al., 2009). Primary antibodies: chicken anti-GFP (Aves Lab, cat# GFP-1010) 1:5000; mouse anti-Dlg (DSHB, 4F3) 1:50; rabbit anti-aPKC (Santa Cruz, Sc-216) 1:1000. Secondary antibodies: Cy2-, Cy3 or Cy5-conjugated goat anti-rabbit IgG, anti-mouse IgG, and anti-chicken IgG (The Jackson ImmunoResearch Lab, 111-225-003, 115-165-003, and 106-175-003), all at 1:400. Images were collected on Olympus FV1000 confocal microscopes (Center for Biologic Imaging, University of Pittsburgh Medical School) and processed in Adobe Photoshop for compositions.

### Image Processing and Quantification

Time-lapse movies were first stabilized by HyperStackReg plug-in in ImageJ. Images or movies containing excessive noisy channels were denoised by PureDenoise plugin in ImageJ prior to quantification. PM localization of GFP or RFP in images or movies were measured in Image J by custom macro scripts. In each image or the first frame of the movie, ROIs approximately 20-40µm^2^ were drawn across selected cell junctions. In most cases, custom macros was used to automatically generate PM masks by threshold-segmentation that was based on the mean pixel value of the ROI. Custom macros were then used to automatically measure PM and cytosolic intensities of each fluorescent protein in ROIs in an image or throughout all the frames of a movie. Backgrounds were manually measured based on the minimal pixel value of the whole image or the first frame of the movie. The PM localization index for each fluorescent protein was auto-calculated by the macro as the ratio of [PM - background]/[cytosol - background]. In live imaging experiments, “Normalized PM Index” was calculated by normalizing (PM Index −1) over the period of recording against the (PM Index −1) at 0 minute. Data were further processed in Excel, visualized and analyzed in Graphpad Prism.

### Tumorigenesis assays

RNAi-resistance wild type or mutant *ubi-dlg::GFP* (“*dlg*^*R*^*\*\*::GFP*”) transgenic alleles were crossed with *ey-Gal4 UAS-Ras*^*V12*^*/CyO-Gal80; UAS-dlg-RNAi/TM6B* and crosses were transferred to a new viral every 3-4 days. F1 progenies from each cross were scored into two groups based on their genotypes: *dlg*^*R*^*\*\*::GFP /ey-Gal UAS-Ras*^*V12*^; *UAS-dlg-RNAi/+* (group#1) or *dlg*^*R*^*\*\*::GFP /ey-Gal UAS-Ras*^*V12*^; *TM6B/+* (group#2). The ratio between groups #1 and #2 was calculated as “Rescue” index for each *dlg*^*R*^*\*\*::GFP* allele. Representative eyes of flies from group#1 were imaged.

