## Supplementary material for "Electrostatic Plasma Membrane Targeting Contributes to Dlg Function in Cell Polarity and Tumorigenesis": Dlg_sup_20201207.pdf

### SUPPLEMNTARY INFORMATION

#### SUPPLEMNTARY TABLES:

##### Table S1. *dlg::GFP* Transgenic Alleles.

| Construct | Vector | Description |
|---|---|---|
| Dlg::GFP | pGU::GFP | Full length Dlg inserted into pGU::GFP |
| Dlg-KR6Q::GFP | pGU::GFP | Dlg::GFP with 6 Lys/Arg residues in PB mutated to Gln |
| Dlg-KR15Q::GFP | pGU::GFP | Dlg::GFP with 15 Lys/Arg residues in PB mutated to Gln |
| Dlg-ΔPDZ::GFP | pGU::GFP | aa1-611 (PDZ domain) deleted from Dlg::GFP |
| Dlg-ΔPDZ-KR6Q::GFP | pGU::GFP | carrying KR6Q mutation in DlgΔPDZ::GFP |
| Dlg-ΔPDZ-KR15Q::GFP | pGU::GFP | carrying KR15Q mutation in DlgΔPDZ::GFP |
| GST-PB | pGEX-4T | Dlg-PB from Dlg::GFP (aa669-722) inserted into pGEX |
| GST-PB-KR6Q | pGEX-4T | Dlg-PB-KR6Q from DlgKR6Q::GFP (aa669-722) inserted into pGEX |
| GST-PB-KR15Q | pGEX-4T | Dlg-PB-KR15Q from DlgKR15Q::GFP (aa669-722) inserted into pGEX |
| Dlg-ΔGUK::GFP | pGU::GFP | aa768-983 (GUK domain) deleted from Dlg::GFP |
| Dlg-S4A::GFP | pGU::GFP | Phospho-serine residues 670, 684, 702, 713 mutated to Ala in Dlg::GFP |
| Dlg-ΔPDZ-S4A::GFP | pGU::GFP | aa1-611 (PDZ domain) deleted from Dlg-S4A::GFP |
| Dlg-S4A-ΔGUK::GFP | pGU::GFP | aa768-983 (GUK domain) deleted from Dlg-S4A::GFP |
| Dlg-m30::GFP | pGU::GFP | aa640 mutated from Leu to Pro in Dlg::GFP |
| Dlg-m30-ΔPDZ::GFP | pGU::GFP | aa1-611 (PDZ domain) deleted from Dlg-m30::GFP |
| Dlg-m30-ΔGUK::GFP | pGU::GFP | aa768-983 (GUK domain) deleted from Dlg-m30::GFP |
| $Dlg^{R}$::GFP | pGU::GFP | TAAGCTGCATGTG AAGCGAAA was converted to TAAACTCCACGTC AAACGCAAG from Dlg::GFP |
| Dlg-$KR6Q^{R}$::GFP | pGU::GFP | TAAGCTGCATGTG AAGCGAAA was converted to TAAACTCCACGTC AAACGCAAG from Dlg-KR6Q::GFP |
| Dlg-$\Delta GUK^{R}$::GFP | pGU::GFP | TAAGCTGCATGTG AAGCGAAA was converted to TAAACTCCACGTC AAACGCAAG from DlgΔGUK::GFP |
| Dlg-$S4A^{R}$::GFP | pGU::GFP | TAAGCTGCATGTG AAGCGAAA was converted to TAAACTCCACGTC AAACGCAAG from DlgS4A::GFP |
| Dlg-$m30^{R}$::GFP | pGU::GFP | TAAGCTGCATGTG AAGCGAAA was converted to TAAACTCCACGTC AAACGCAAG from Dlgm30::GFP |

SUPPLEMENTARY FIGURES:

### Figure S1

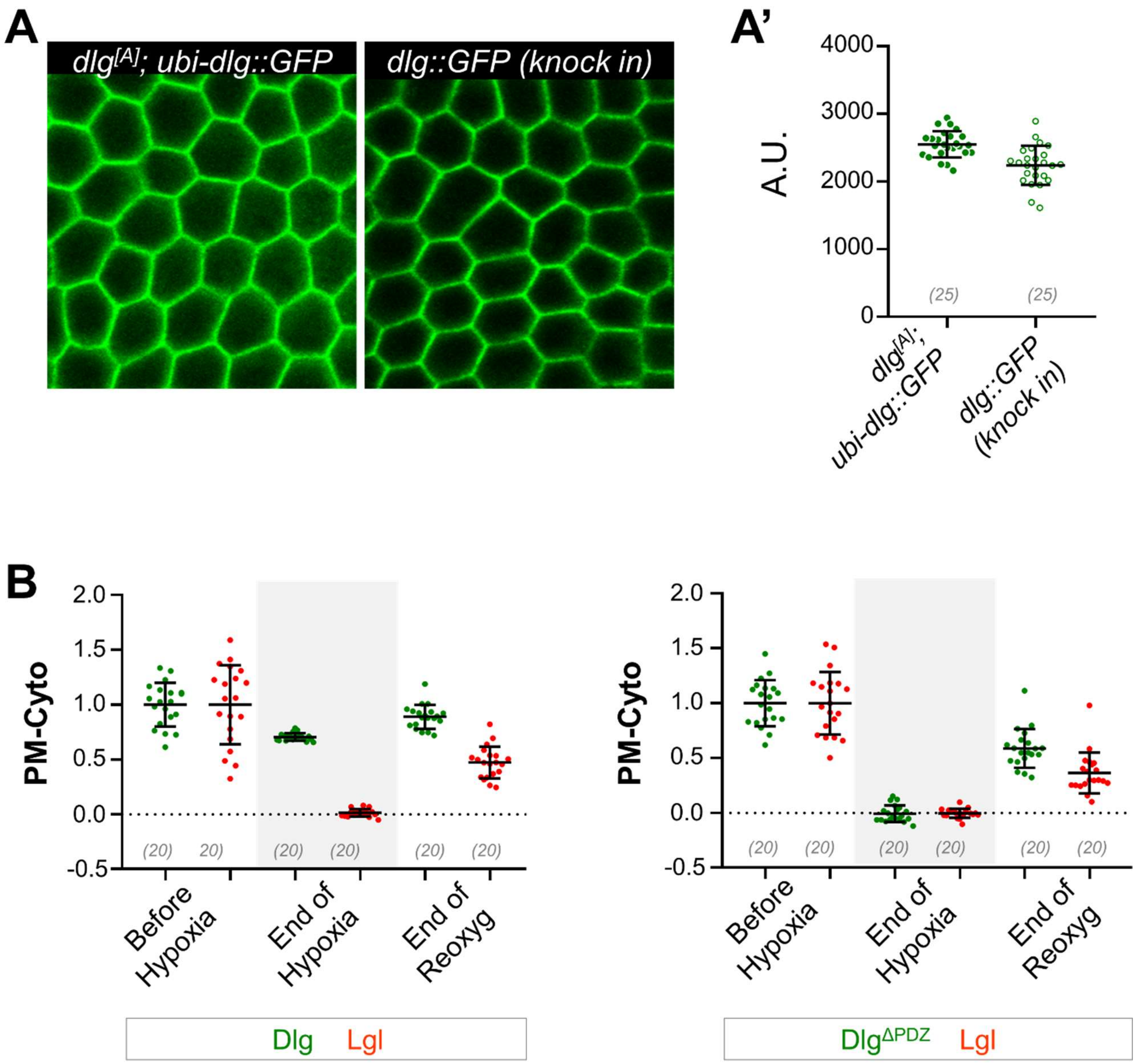


**Figure S1. Overexpressed ubi-Dlg::GFP and endogenous Dlg::GFP$^{KI}$ show similar expression levels.**

(**A**) Follicular cells in ovaries at similar stages from rescued *dlg[A] ubi-dlg::GFP* females and from *dlg::GFP$^{KI}$* knock-in females. Ovaries were mounted at the same slide and images were captured under identical acquisition parameters. (**A'**) Quantifications of ubi-

Dlg::GFP and Dlg::GFP$^{KI}$, measured by the average intensity of ROIs over multiple cells. A.U.: arbitrary unit.

(**B**) Quantifications of PM levels of Dlg$^{KI}$::GFP and Dlg$^{\Delta PDZ}$::GFP in Movies S1 and S3 (also shown in Fig. 1D,E) at the time points of before hypoxia, end of hypoxia, and end of reoxygenation ("reoxyg"), respectively. Lgl::RFP serves as controls. "PM-Cyto": (PM intensity - cytosolic intensity) at each ROIs that is normalized to the average value at the time point before hypoxia. ROI sets are the same as used for the quantification in Figure 1D', E'.

In parentheses: sample numbers.

### Figure S2

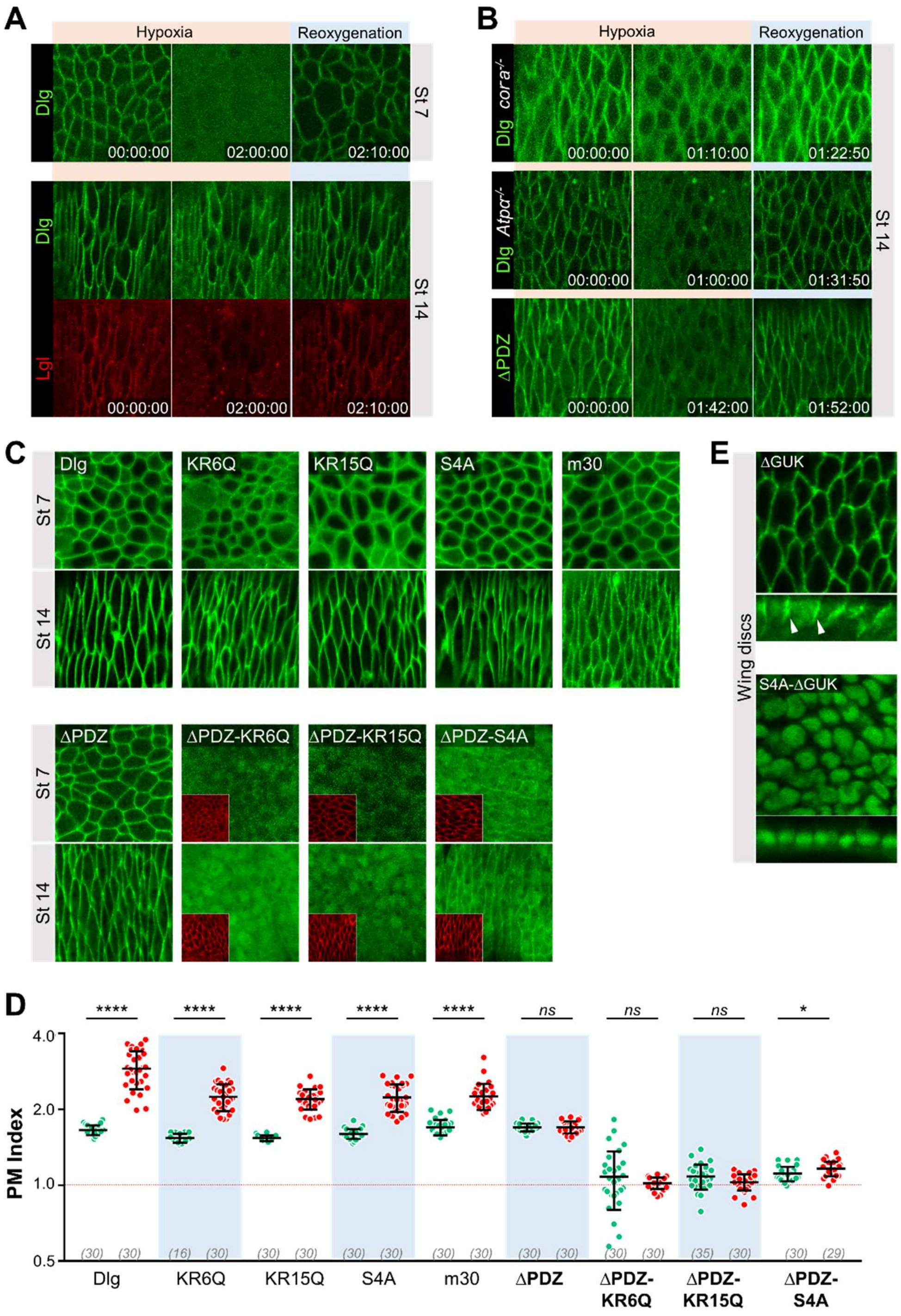


**Figure S2. Septate junctions retain Dlg PM localization in redundancy to electrostatic PM targeting.**

(**A**) PM localization of Dlg::GFP in early (stage 7, “St 7”) but not late (stage 14, “St 14”) embryonic epithelia was sensitive to hypoxia. Stage 14 embryo was also expressing Lgl::mCherry serving as a positive control for hypoxia treatment.

(**B**) PM localization of $Dlg^{KI}$::GFP in embryonic epithelial cells was sensitive to hypoxia in late stage embryos of *cora*$^{-/-}$ or *Atpα*$^{-/-}$. PM localization of $Dlg^{\Delta PDZ}$::GFP in late stage wild type embryo was also sensitive to hypoxia.

(**C**) PM localization of wild type and mutant Dlg::GFP in early and late embryonic epithelial cells.  Embryos expressing $Dlg^{\Delta PDZ\text{-}KR6Q}$::GFP, $Dlg^{\Delta PDZ\text{-}KR15Q}$::GFP and $Dlg^{\Delta PDZ\text{-}S4A::GFP}$, were also expressing Lgl::mCherry (insets).

(**D**) Quantifications of PM localizations of wild type and mutant Dlg::GFP in **C**. For each mutant, green and red dots represent the PM quantification data from early St7 and late St14 embryos, respectively.

(**E**) Subcellular localization of $Dlg^{\Delta GUK}$::GFP and $Dlg^{S4A\text{-}\Delta GUK}$::GFP in wing disc epithelial cells. White arrowheads in cross section view of $Dlg^{\Delta GUK}$::GFP indicate the localization to SJ, which is not seen in the cross section view of $Dlg^{S4A\text{-}\Delta GUK}$::GFP.

Time stamps in A and B are in *hh:mm:ss* format. ****: $p$<0.00001; *: $p$<0.001; *ns*: $p$>0.05.

In parentheses: sample numbers.

### Figure S3

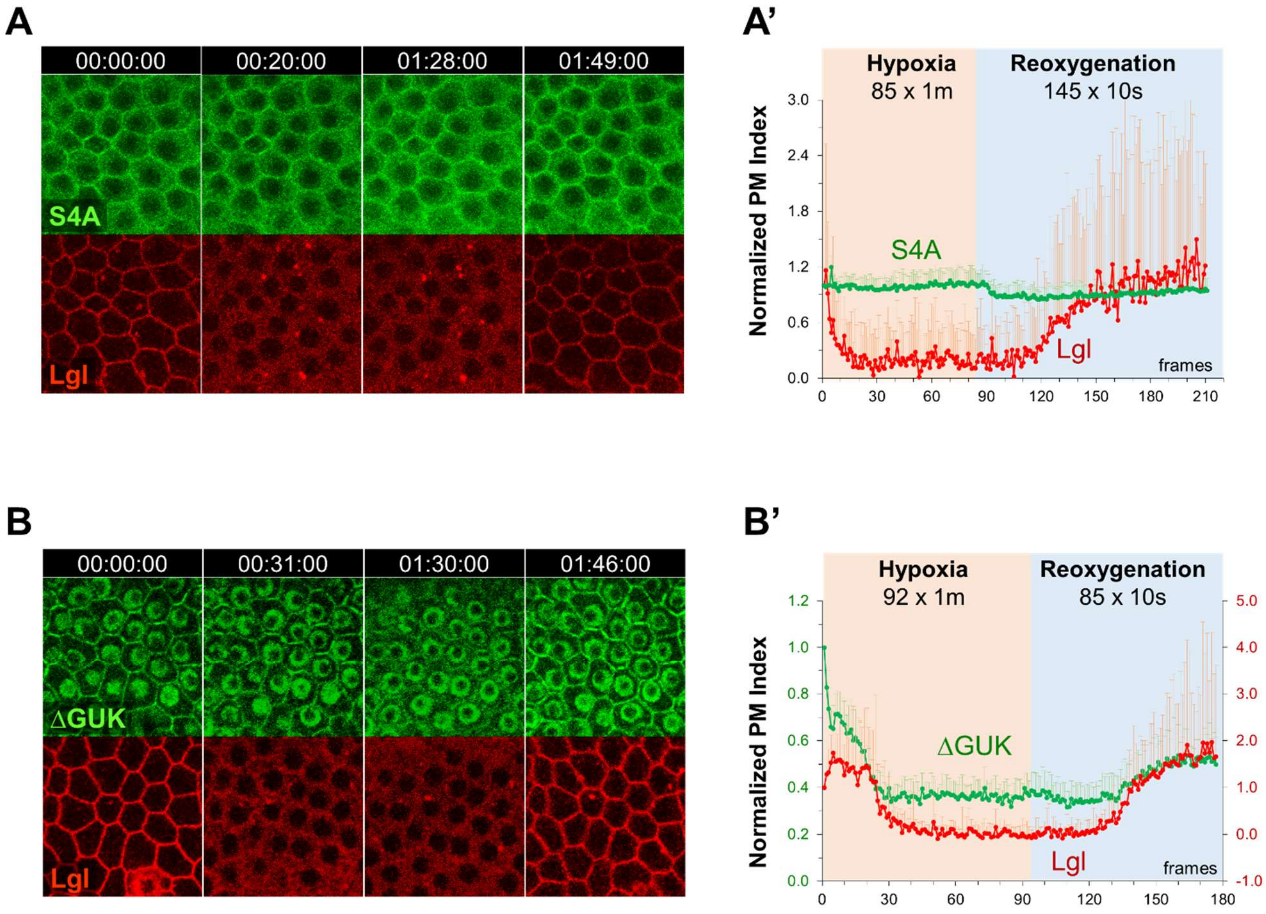


**Figure S3. PM targeting of Dlg$^{\Delta GUK}$::GFP but not Dlg$^{S4A}$::GFP is sensitive to hypoxia.**

(**A, B**) Selected frames of time-lapse recordings of *lgl::mCherry dlg$^{S4A}$::GFP* (B, Movie S6) or *lgl::mCherry dlg$^{\Delta GUK}$::GFP* (A, Movie S7) follicular cells undergoing hypoxia and reoxygenation.

(**A', B'**) Quantification of PM localizations of Lgl::mCherry and Dlg$^{S4A}$::GFP in A (A', *n*=24) and Lgl::mCherry and Dlg$^{\Delta GUK}$::GFP (B', *n*=15).

Time stamps in *hh:mm:ss* format.

### Figure S4

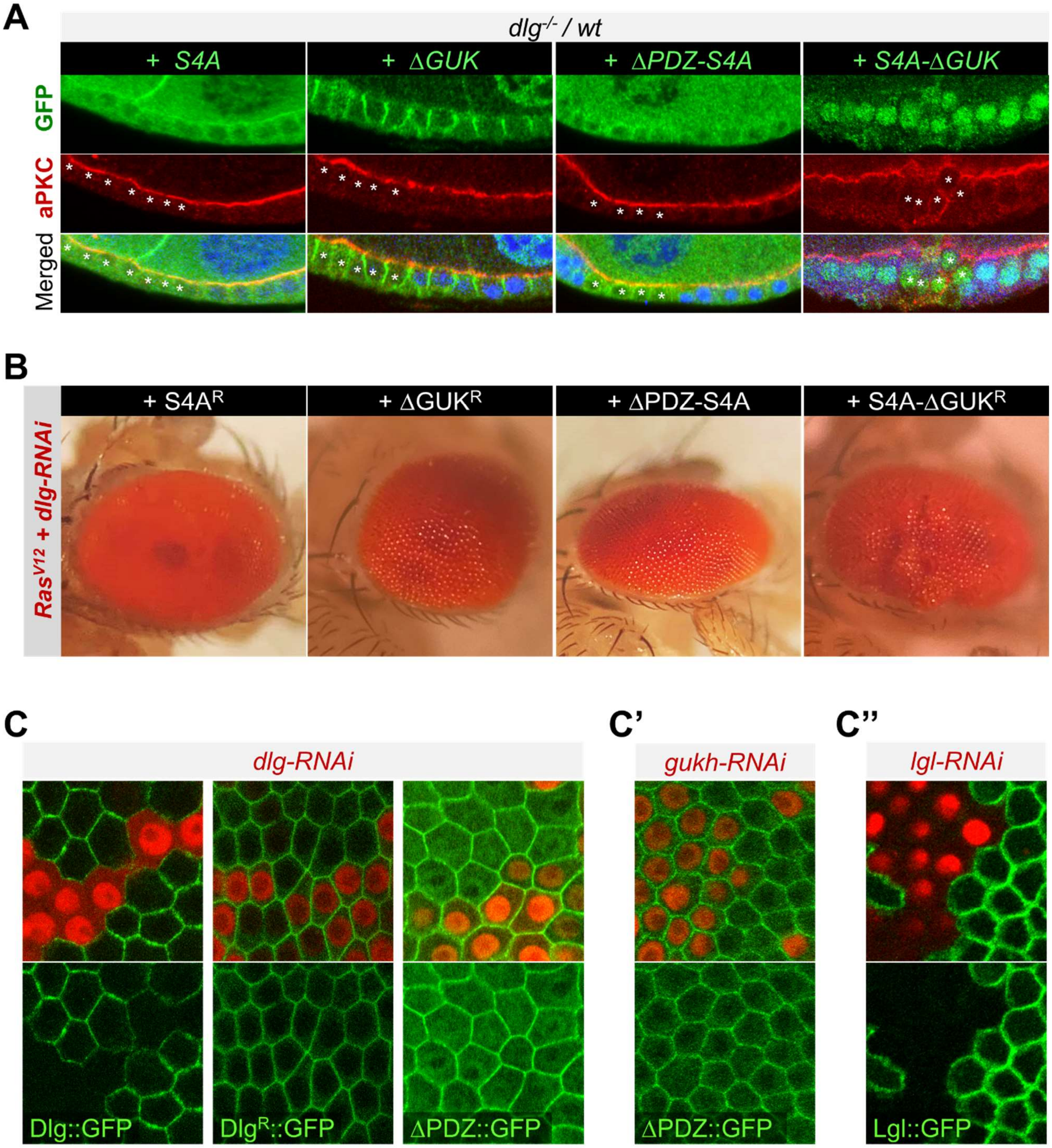


**Figure S4. Electrostatic PM targeting of Dlg regulates cell polarity and**

**tumorigenesis.**

(**A**) Representative immunostaining images of follicular cells showing the potential rescue of *dlg*$^{-/-}$ clones (marked by the loss of RFP, colored in blue in all merged images) by *dlg*$^{S4A}$*::GFP* ("*S4A*"), *dlg*$^{\Delta GUK}$*::GFP* ("*ΔGUK*"), *dlg*$^{\Delta PDZ\text{-}S4A}$*::GFP* ("*ΔPDZ-S4A*") and *dlg*$^{S4A\text{-}\Delta GUK}$*::GFP* ("*S4A-ΔGUK*"), respectively. All samples were stained with anti-GFP (green) and anit-aPKC (red) antibodies. Asterisks highlight *dlg*$^{-/-}$ mutant cells.

(**B**) Representative images of eyes from adult flies of *ey-Gal4, UAS-Ras*$^{V12}$*, UAS-dlg-RNAi* ("*Ras*$^{V12}$ + *dlg-RNAi*") in combination with additional expression of RNAi-resistant *dlg*$^{R\text{-}S4A}$*::GFP* ("*S4A*$^{R}$"), *dlg*$^{R\text{-}\Delta GUK}$*::GFP* ("*ΔGUK*$^{R}$"), *dlg*$^{\Delta PDZ\text{-}S4A}$*::GFP* ("*ΔPDZ-S4A*") and *dlg*$^{R\text{-}S4A\text{-}\Delta GUK}$*::GFP* ("*S4A-ΔGUK*$^{R}$"), respectively.

**(C)** Dlg::GFP, but not Dlg$^{R}$::GFP or Dlg$^{\Delta PDZ}$::GFP, was efficiently knocked down in *dlg-RNAi* follicular cells (labeled by nuclear RFP).

**(C')** PM localization of Dlg$^{\Delta PDZ}$::GFD was not affected in *gukh-RNAi* follicular cells (labeled by nuclear RFP).

**(C'')** Lgl::GFP was efficiently knocked down in *lgl-RNAi* follicular cells (labeled by nuclear RFP).

#### SUPPLEMENTARY MOVIES

##### Movie S1. Acute and reversible loss of PM localization of Dlg::GFP$^{KI}$ and Lgl::mCherry under hypoxia in follicular cells

Ovaries from a 2-day old *dlg::GFP*$^{KI}$*, lgl::mCherry* female were dissected and imaged live in an environment-controlled micro chamber. Hypoxic (0.5% $O_2$) gas was flashed into the chamber at 0 minute to induce hypoxia and normal air was flashed into chamber at 111 minutes for reoxygenation. Time intervals are 3 minutes during hypoxia and 10 seconds during reoxygenation. Note the incomplete loss of Dlg::GFP from the PM at the end of hypoxia. Time stamp: *hh:mm:ss*.

##### Movie S2. Acute and reversible loss of PM localization of ubi-Dlg::GFP under hypoxia in follicular cells

Ovaries from a 2-day old *ubi-dlg::GFP* female were dissected and imaged live similarly as in Movie S1. Reoxygenation started at 110 minutes in the movie. Time intervals are 5 minute during hypoxia and 30 seconds during reoxygenation. Time stamp: *hh:mm:ss*.

**Movie S3. Acute and reversible loss of PM localization of Dlg$^{\Delta PDZ}$::GFP and Lgl::mCherry under hypoxia**

Ovaries from a 2-day old *lgl::mCherry*, *dlg$^{\Delta PDZ}$::GFP* female were dissected and imaged live similarly as in Movie S1. Reoxygenation started at 36 minutes in the movie. Time intervals are 1 minute during hypoxia and 10 seconds during reoxygenation. Time stamp: *hh:mm:ss*.

**Movie S4. PM localization of Dlg::GFP in *PI4KIIIα-RNAi* cells showed accelerated loss under hypoxia and delayed recovery under reoxygenation**

*dlg$^{KI}$::GFP, PI4KIIIα-RNAi* females were heat-shocked and ovaries were dissected three days after and imaged live similarly as in Movie S1. Reoxygenation starts at 96 minutes in the movie. Time intervals are 3 minutes during hypoxia and 10 seconds during reoxygenation. *PI4KIIIα-RNAi* cells were labelled by RFP. Time stamp: *hh:mm:ss*.

**Movie S5. PM localization of Dlg$^{\Delta PDZ}$::GFP in *PI4KIIIα-RNAi* cells showed accelerated loss under hypoxia and delayed recovery under reoxygenation**

*dlg$^{\Delta PDZ}$::GFP, PI4KIIIα-RNAi* females were heat-shocked and ovaries were dissected three days after and imaged live similarly as in Movie S1. Reoxygenation starts at 20 minutes in the movie. Time intervals are 1 minute during hypoxia and 10 seconds during reoxygenation. *PI4KIIIα-RNAi* cells were labelled by RFP. Time stamp: *hh:mm:ss*.

**Movie S6. PM localization of Dlg$^{S4A}$::GFP is resistant to hypoxia**

Ovaries from a 2-day old *lgl::mCherry dlg$^{S4A}$::GFP* female were dissected and imaged live similarly as in Movie S1. Reoxygenation starts at 85 minutes in the movie. Time intervals are 1 minute during hypoxia and 10 seconds during reoxygenation. Time stamp: *hh:mm:ss*.

**Movie S7. Acute and reversible loss of PM localization of Dlg$^{\Delta GUK}$::GFP under hypoxia**

Ovaries from a 2-day old *dlg$^{\Delta GUK}$::GFP; lgl::mCherry* female were dissected and imaged live similarly as in Movie S1. Reoxygenation starts at 92 minutes in the movie. Time intervals are 1 minute during hypoxia and 10 seconds during reoxygenation. Time stamp: *hh:mm:ss*.

**Genotypes of *Drosophila* Samples in Figures.**

**Figure 1:**

**(C)** *w; ubi-dlg::GFP/CyO*

*w; ubi-dlg$^{KR6Q}$::GFP/CyO*

*w; ubi-dlg$^{KR15Q}$::GFP/CyO*

*w; ubi-dlg$^{\Delta PDZ}$::GFP/CyO*

*w; ubi-dlg$^{\Delta PDZ\text{-}KR6Q}$::GFP/CyO*

*w; ubi-dlg$^{\Delta PDZ\text{-}KR15Q}$::GFP/CyO*

**(D)** *w dlg::GFP$^{KI}$/+; lgl::mCherry/+*

**(E)** *w; ubi-dlg$^{\Delta PDZ}$::GFP/lgl::mCherry*

**Figure 2:**

**(A)** *w UAS-mRFP-FKBP-5'Ptas/dlg$^{KI}$::GFP;; hs-FLP Act5C(FRT.CD2)-Gal4 UAS-RFP$^{NLS}$/ UAS-Lck-FRB::CFP*

*w UAS-mRFP-FKBP-5'Ptas/+; ubi-dlg$^{\Delta PDZ}$::GFP/+; hs-FLP Act5C(FRT.CD2)-Gal4 UAS-RFP$^{NLS}$/UAS-Lck-FRB::CFP*

**(B)** *w dlg::GFP$^{KI}$;; hs-FLP Act5C(FRT.CD2)-Gal4 UAS-RFP$^{NLS}$/UAS-PI4KIIIα-RNAi*

*w; ubi-dlg$^{\Delta PDZ}$::GFP; hs-FLP Act5C(FRT.CD2)-Gal4 UAS-RFP$^{NLS}$/UAS-PI4KIIIα-RNAi*

**Figure 3:**

**(A)**

*w; ubi-dlg::GFP/CyO*

*w; ubi-dlg$^{\Delta PDZ}$::GFP/CyO*

*w; ubi-dlg$^{\Delta GUK}$::GFP/CyO*

*w; ubi-dlg$^{S4A}$::GFP/CyO*

*w; ubi-dlg$^{S4D}$::GFP/CyO*

*w; ubi-dlg*$^{\Delta PDZ\text{-}S4A}$*::GFP/CyO*

*w; ubi-dlg*$^{S4A\text{-}\Delta GUK}$*::GFP/CyO*

*w; ubi-dlg*$^{m30}$*::GFP/CyO*

*w; ubi-dlg*$^{m30\text{-}\Delta PDZ}$*::GFP/CyO*

*w;ubi-dlg*$^{m30\text{-}\Delta GUK}$*::GFP/CyO*

**(B)** *w dlg::GFP*$^{KI}$*/+;; hs-FLP Act5C(FRT.CD2)-Gal4 UAS-RFP*$^{NLS}$*/ UAS-scrib-RNAi*

*w; ubi-dlg*$^{\Delta PDZ}$*::GFP/+; hs-FLP Act5C(FRT.CD2)-Gal4 UAS-RFP*$^{NLS}$*/ UAS-scrib-RNAi*

*w; ubi-dlg*$^{KR6Q}$*::GFP/+; hs-FLP Act5C(FRT.CD2)-Gal4 UAS-RFP*$^{NLS}$*/ UAS-scrib-RNAi*

*w; ubi-dlg*$^{S4A}$*::GFP/+; hs-FLP Act5C(FRT.CD2)-Gal4 UAS-RFP*$^{NLS}$*/ UAS-scrib-RNAi*

*w; ubi-dlg*$^{\Delta GUK}$*::GFP/+; hs-FLP Act5C(FRT.CD2)-Gal4 UAS-RFP*$^{NLS}$*/ UAS-scrib-RNAi*

*w; ubi-dlg*$^{m30}$*::GFP/+; hs-FLP Act5C(FRT.CD2)-Gal4 UAS-RFP*$^{NLS}$*/ UAS-scrib-RNAi*

**(C)** *w dlg::GFP*$^{KI}$*/+; ; hs-FLP Act5C(FRT.CD2)-Gal4 UAS-RFP*$^{NLS}$*/ UAS-lgl-RNAi*

*w; ubi-dlg*$^{\Delta PDZ}$*::GFP/+; hs-FLP Act5C(FRT.CD2)-Gal4 UAS-RFP*$^{NLS}$*/ UAS-lgl-RNAi*

*w; ubi-dlg*$^{KR6Q}$*::GFP/+; hs-FLP Act5C(FRT.CD2)-Gal4 UAS-RFP*$^{NLS}$*/ UAS-lgl-RNAi*

*w; ubi-dlg*$^{S4A}$*::GFP/+; hs-FLP Act5C(FRT.CD2)-Gal4 UAS-RFP*$^{NLS}$*/ UAS-lgl-RNAi*

*w; ubi-dlg*$^{\Delta GUK}$*::GFP/+; hs-FLP Act5C(FRT.CD2)-Gal4 UAS-RFP*$^{NLS}$*/ UAS-lgl-RNAi*

*w; ubi-dlg*$^{m30}$*::GFP/+; hs-FLP Act5C(FRT.CD2)-Gal4 UAS-RFP*$^{NLS}$*/ UAS-lgl-RNAi*

**Figure 4:**

**(A)** *w dlg*$^{[A]}$ *FRT*$^{19A}$*/ubi-RFP FRT*$^{19A}$

*w dlg*$^{[A]}$ *FRT*$^{19A}$*/ubi-RFP FRT*$^{19A}$*; ubi-dlg::GFP/+*

*w dlg*$^{[A]}$ *FRT*$^{19A}$*/ubi-RFP FRT*$^{19A}$*; ubi-dlg*$^{\Delta PDZ}$*::GFP/+*

*w dlg*$^{[A]}$ *FRT*$^{19A}$*/ubi-RFP FRT*$^{19A}$*; ubi-dlg*$^{KR6Q}$*::GFP/+*

*w dlg*$^{[A]}$ *FRT*$^{19A}$*/ubi-RFP FRT*$^{19A}$*; ubi-dlg*$^{\Delta PDZ\text{-}KR6Q}$*::GFP/+*

*w dlg*$^{[A]}$ *FRT*$^{19A}$*/ubi-RFP FRT*$^{19A}$*; ubi-dlg*$^{S4A}$*::GFP/+*

*w dlg*$^{[A]}$ *FRT*$^{19A}$*/ubi-RFP FRT*$^{19A}$*; ubi-dlg*$^{m30}$*::GFP/+*

*w dlg*$^{[A]}$ *FRT*$^{19A}$*/ubi-RFP FRT*$^{19A}$ *ubi-dlg*$^{m30\text{-}\Delta PDZ}$*::GFP/+*

*w dlg$^{[A]}$ FRT$^{19A}$/ubi-RFP FRT$^{19A}$; ubi-dlg$^{m30\text{-}\Delta GUK}$::GFP/+*

**(B)** Top row:

*w; ey-Gal4 UAS-Ras$^{V12}$/+*

*w; ey-Gal4 UAS-Ras$^{v12}$/+; UAS-dlg-RNAi/+*

*w; ey-Gal4 UAS-Ras$^{V12}$/ubi-dlg$^{R}$::GFP; UAS-dlg-RNAi/+*

*w; ey-Gal4 UAS-Ras$^{V12}$/ubi-dlg$^{\Delta PDZ}$::GFP; UAS-dlg-RNAi/+*

*w; ey-Gal4 UAS-Ras$^{V12}$/ubi-dlg$^{KR6Q\text{-}R}$::GFP; UAS-dlg-RNAi/+*

Bottom row:

*w; ey-Gal4/+; UAS-dlg-RNAi/+*

*w; ey-Gal4 UAS-Ras$^{V12}$/ubi-dlg$^{\Delta PDZ\text{-}KR6Q}$::GFP; UAS-dlg-RNAi/+*

*w; ey-Gal4 UAS-Ras$^{V12}$/ubi-dlg$^{m30\text{-}R}$::GFP; UAS-dlg-RNAi/+*

*w; ey-Gal4 UAS-Ras$^{V12}$/ubi-dlg$^{m30\text{-}\Delta PDZ}$::GFP; UAS-dlg-RNAi/+*

*w; ey-Gal4 UAS-Ras$^{V12}$/ubi-dlg$^{m30\text{-}\Delta GUK\text{-}R}$::GFP; UAS-dlg-RNAi/+*

**Figure S1:**

**(A)** *w dlg$^{[A]}$ FRT$^{19A}$/ubi-RFP FRT$^{19A}$; ubi-dlg::GFP/ ubi-dlg::GFP*

*dlg::GFP$^{KI}$/ dlg::GFP$^{KI}$*

**Figure S2:**

**(A)** *w*; *ubi-dlg::GFP/ubi-dlg::GFP*

*w; dlg::GFP$^{KI}$/Lgl::mCherry*

**(B)** *w dlg::GFP$^{KI}$/+; cora$^{5}$/ cora$^{5}$*

*w dlg::GFP$^{KI}$/+; Atpα$^{DTS2A3}$/ Atpα$^{DTS2A3}$*

*w; ubi-dlg$^{\Delta PDZ}$::GFP /CyO*

**(C)** Top row:

*w; ubi-dlg::GFP/CyO*

*w; ubi-dlg*$^{KR6Q}$*::GFP /CyO*

*w;ubi-dlg*$^{KR15Q}$*::GFP /CyO*

*w; ubi-dlg*$^{S4A}$*::GFP /CyO*

*w; ubi-dlg*$^{m30}$*::GFP /CyO*

Bottom row

*w; ubi-dlg*$^{\Delta PDZ}$*::GFP/CyO*

*w; ubi-dlg*$^{\Delta PDZ\text{-}KR6Q}$*::GFP / lgl::mCherry*

*w; ubi-dlg*$^{\Delta PDZ\text{-}KR15Q}$*::GFP / lgl::mCherry*

*w; ubi-dlg*$^{\Delta PDZ\text{-}S4A}$*::GFP / lgl::mCherry*

**(E)** *w; ubi-dlg*$^{\Delta GUK}$*::GFP/CyO*

*w; ubi-dlg*$^{S4A\text{-}\Delta GUK}$*::GFP/CyO*

**Figure S3:**

**(A)** *w; ubi-dlg*$^{\Delta GUK}$*::GFP / lgl::mCherry*

**(B)** *w; ubi-dlg*$^{S4A}$*::GFP / lgl::mCherry*

**Figure S4:**

**(A)** *w dlg*$^{[A]}$ *FRT*$^{19A}$*/ubi-RFP FRT*$^{19A}$*; ubi-dlg*$^{S4A}$*::GFP/+*

*w dlg*$^{[A]}$ *FRT*$^{19A}$*/ubi-RFP FRT*$^{19A}$*; ubi-dlg*$^{\Delta GUK}$*::GFP/+*

*w dlg*$^{[A]}$ *FRT*$^{19A}$*/ubi-RFP FRT*$^{19A}$*; ubi-dlg*$^{\Delta PDZ\text{-}S4A}$*::GFP/+*

*w dlg*$^{[A]}$ *FRT*$^{19A}$*/ubi-RFP FRT*$^{19A}$*; ubi-dlg*$^{S4A\ \text{-}\Delta GUK}$*::GFP/+*

**(B)** *w; ey-Gal4 UAS-Ras*$^{V12}$*/ubi-dlg*$^{S4A\text{-}R}$*::GFP; UAS-dlg-RNAi/+*

*w; ey-Gal4 UAS-Ras*$^{V12}$*/ubi-dlg*$^{\Delta GUK\text{-}R}$*::GFP; UAS-dlg-RNAi/+*

*w; ey-Gal4 UAS-Ras*$^{V12}$*/ubi-dlg*$^{\Delta PDZ\text{-}S4A}$*::GFP; UAS-dlg-RNAi/+*

*w; ey-Gal4 UAS-Ras*$^{V12}$*/ubi-dlg*$^{S4A\text{-}\Delta GUK\text{-}R}$*::GFP; UAS-dlg-RNAi/+*

**(C)** *w dlg::GFP/+; UAS-dlg-RNAi/+; hs-FLP Act5C(FRT.CD2)-Gal4 UAS-RFP*$^{NLS}$ */+*

*w; UAS-dlg-RNAi /ubi-dlg*$^{R}$*-GFP; hs-FLP Act5C(FRT.CD2)-Gal4 UAS-RFP*$^{NLS}$ */+*

*w; dlg-RNAi /ubi-dlg*$^{\Delta PDZ}$*-GFP; hs-FLP Act5C(FRT.CD2)-Gal4 UAS-RFP*$^{NLS}$ */+*

**(C')** *w dlg::GFP/+; UAS-gukh-RNAi/+; hs-FLP Act5C(FRT.CD2)-Gal4 UAS-RFP*$^{NLS}$ */+*

**(C'')** *w; lgl::GFP hs-FLP/ lgl-RNAi; Act5C(FRT.CD2)-Gal4 UAS-RFP /+*
